# Atg39 selectively captures inner nuclear membrane into lumenal vesicles for delivery to the autophagosome

**DOI:** 10.1101/2021.02.22.432332

**Authors:** Sunandini Chandra, Philip J. Mannino, David J. Thaller, Nicholas R. Ader, Megan C. King, Thomas J. Melia, C. Patrick Lusk

## Abstract

Mechanisms that turnover components of the nucleus and inner nuclear membrane (INM) remain to be fully defined. We explore how components of the INM are selected by a cytosolic autophagy apparatus through a transmembrane nuclear envelope-localized cargo adaptor, Atg39. A split-GFP reporter shows that Atg39 localizes to the outer nuclear membrane (ONM) and thus targets the INM across the nuclear envelope lumen. Consistent with this, sequence elements that confer both nuclear envelope localization and a membrane remodeling activity are mapped to the Atg39 lumenal domain; these lumenal motifs are required for the autophagy-mediated degradation of an integral INM protein. Interestingly, correlative light and electron tomography shows that the overexpression of Atg39 leads to the expansion of the ONM and the enclosure of a network of INM-derived vesicles in the nuclear envelope lumen. Thus, we propose an outside-in model of nucleophagy where INM is delivered into vesicles in the nuclear envelope lumen, which can be targeted by the autophagosome.

## Introduction

The function of the nuclear envelope (NE) is conferred by its biochemical constituents that populate an inner nuclear membrane (INM) with peripherally associated nuclear lamina, an outer nuclear membrane (ONM), and a nuclear pore membrane^1^. The latter defines connections between the INM and ONM where embedded nuclear pore complexes (NPCs) control molecular traffic between the nucleus and cytoplasm^2, 3^. Although we have a considerable understanding of the mechanisms that underly molecular exchange through NPCs, it is less well understood how the NE proteome is turned over under either physiological or pathological conditions.

The need to clear damaged or defective proteins from the nucleus and NE is underscored by the accumulation of nuclear protein aggregates in several human diseases^4^. Further, both NPCs^5^ and the nuclear lamins accumulate damage with age^6^ and defects in nuclear transport^7–9^ and NPC injury may be a cause of certain forms of neurodegeneration including amyotrophic lateral sclerosis^10^. Interestingly, the protein constituents of NPCs are also characterized by long half lives in neurons^11–13^, which may indicate that they are challenging to productively turnover. Indeed, it is hard to conceptualize how cells might remove these massive macromolecular assemblies without compromising NE integrity. Nonetheless, there is evidence that NPCs may be excised from the NE in both metazoan^14^ and in yeast model systems^15–17^. While the endosomal sorting complexes required for transport (ESCRT)^14, 16^ and the macroautophagy^16, 17^ machinery have been implicated in these events, the molecular and morphological steps in these pathways are just beginning to come to light.

Like NPCs, there is evidence that the lamins can be turned over with several molecular links implicating macroautophagy in this process^18–21^. Macroautophagy (hereafter called “autophagy”) is a catabolic mechanism that delivers protein aggregates, lipids and parts of (and in some cases, whole) organelles to lysosomes for degradation^22^. It begins with the formation of a phagophore membrane that is defined by the covalent coupling of the ubiquitin-like protein LC3 (Atg8, in yeast) directly to phosphatidylethanolamine^23^. The phagophore expands around the cargo, ultimately sealing the cargo inside a closed double membrane organelle called the autophagosome^24^. The autophagosome ultimately fuses with lysosomes (or vacuoles, in yeast) where cargo is degraded^25^.

Interestingly, there is evidence that LC3 can direct a form of nuclear autophagy (nucleophagy) by binding to Lamin B1 in the context of oncogene activation: this interaction plays a part in the selective clearance of Lamin B1 from the INM^18^. However, how a cytosolic phagophore selectively targets the INM or nucleoplasm across the double membraned NE remains unknown. Indeed, in most cases of selective organelle targeting, autophagy cargo adaptors bind to specific proteins and recruit the autophagy machinery to initiate phagophore expansion around themselves^23, 26, 27^. While such a nuclear-specific cargo adaptor has not been identified in metazoans, budding yeast have Atg39 (a.k.a. Esm1^28^), a putative type II transmembrane protein^28^ that localizes at the NE and is required for the autophagic degradation of both INM and nucleoplasmic proteins^17, 29–33^, but not NPCs^16^. Thus, Atg39 is the essential cog in the macroautophagic clearance of INM, but how INM is recognized and delivered to the cytoplasm remains unknown. Here, we explore the mechanism of Atg39-mediated nucleophagy in budding yeast. The data support that Atg39 acts from the ONM and connects to the INM through its lumenal domain. The lumenal domain has functional elements that are required for NE remodeling and the capture of INM cargo into NE-blebs that can be targeted by autophagy. By using correlative light and electron microscopy (CLEM) and tomography and focused ion beam-scanning EM (FIB-SEM) to visualize NE bleb ultrastructure, we observe the capture of INM into vesicles in the NE lumen. We propose a model where nucleophagy proceeds through an outside-in mechanism where putative translumenal interactions coordinate INM and ONM remodeling to ultimately deliver INM cargo to the autophagosome.

## Results

### Atg39 accumulates at the ONM

Key unknowns to unraveling the nucleophagy mechanism are determining whether Atg39 acts from the ONM or the INM (or both), and whether, like other cargo adaptors^34–36^ it has any inherent membrane remodeling activity. To address the former, we took advantage of a recently developed split-GFP reporter system used to catalogue the INM proteome^37^. The system exploits a series of mCherry-tagged reporter proteins that are expressed as fusions to GFP^11^, a 4 kD fragment of GFP, and are localized in the nucleus (GFP^11^-mCherry-Pus1; Fig. 1a) and ER. The two ER reporters differ in that GFP^11^ either faces the lumen (mCherry-Scs2_TM_-GFP^11^) or the cytosol (GFP^11^-mCherry-Scs2TM; Fig. 1a). In these backgrounds, the rest of GFP (GFP^1–10^) was expressed on either the N- or C-terminus of Atg39 (Fig. 1a) from a galactose-inducible (*GAL1*) promoter; the reconstitution of a fluorescent GFP provides evidence of physical proximity with which to infer localization.

**Fig. 1.**
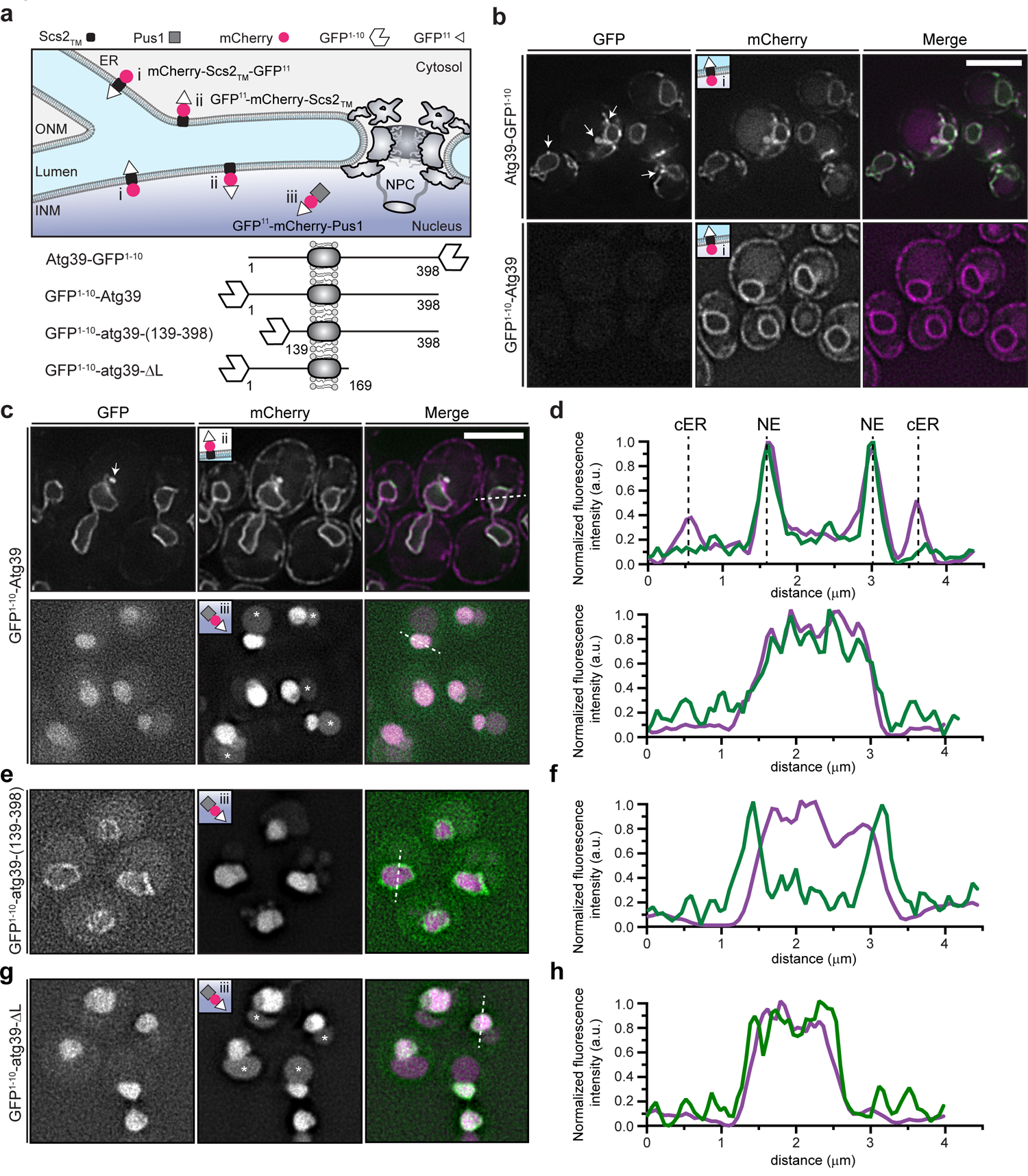
Atg39 accumulates at the ONM. **a**, Schematic of localization and topology of split-GFP constructs. The GFP^11^-mCherry reporter fusion proteins (i, ii, iii) are constructed as shown in key with localization diagrammed. INM and ONM are inner and outer nuclear membrane, respectively. At bottom are schematics of the Atg39 GFP^1–10^ fusions and truncations lacking N or C-termini. The Atg39 transmembrane (TM) domain is depicted as a grey oval. Numbers indicate amino acids. **b**, Deconvolved fluorescence micrographs of cells expressing indicated Atg39 fusion proteins and mCherry-reporters (see inset). GFP, mCherry and merged images shown. Arrowhead points to NE bleb. Scale bar is 5 µm. **c**, Deconvolved fluorescence micrographs of cells expressing GFP^1–10^-Atg39 and indicated mCherry-reporter (inset). GFP, mCherry and merged images shown. Scale bar is 5 µm. Asterisks indicate vacuolar autofluorescence. **d**, Normalized line profiles of GFP (green) and mCherry (magenta) fluorescence (in arbitrary units, a.u.) bisecting cells as indicated by dotted lines in corresponding top and bottom merge panels from **c**. Position of NE (nuclear envelope) and cER (cortical ER) is indicated by dotted lines. **e**, As in **c,** but with cells expressing GFP^1–10^-atg39-(139-398). **f**, As in **d**, but with cells from **e**. **g**, As in **c**, but with cells expressing GFP^1–10^-atg39-ΔL. Asterisks indicate vacuolar autofluorescence. **h**, As in **d**, but with cells from **g**.

Taking advantage of the mCherry-Scs2_TM_-GFP^11^ ER lumenal reporter (Fig. 1a, i), we first confirmed the proposed type II topology^28, 29^ of Atg39 as only Atg39-GFP^1–10^, but not GFP^1–10^-Atg39 resulted in visible GFP fluorescence localized at the NE (Fig. 1b). We also noted that there were NE extensions or “blebs” at sites of reconstituted GFP fluorescence (Fig. 1b, arrows) suggesting that Atg39 expression impacted NE morphology, consistent with prior data^28^. These blebs were also observed when GFP^1–10^-Atg39 was expressed alongside the cytosolic-facing ER reporter (Fig. 1c, top panels, arrow). Consistent with the conclusion that GFP^1–10^-Atg39 is localized specifically to the NE, the ER reporter itself was distributed throughout the NE and ER but the GFP fluorescence was only observed at the nuclear periphery and within the blebs extending from the NE. This was particularly evident when line profiles were drawn that bisected the entire cell and nucleus: only the NE, and not the cortical (c) ER peaks of the mCherry fluorescence (magenta) overlapped with the reconstituted GFP-Atg39 fluorescence (green)(Fig. 1d).

As the ER reporters can access both the ONM and INM (Fig. 1a), we tested whether Atg39 could reach the INM using the nucleoplasmic reporter (Fig. 1a, iii). The extralumenal domain of Atg39 is predicted to be unstructured with a MW of 16 kD. Thus, even with the addition of the GFP^1–10^, it should, in principle, be able to pass through the peripheral channels along the pore membrane, which are thought to restrict passage of extralumenal domains larger than ∼60 kD^37, 38^. Interestingly, although we observed low levels of GFP fluorescence when GFP^1–10^-Atg39 was expressed with the nucleoplasmic reporter (Fig. 1c, bottom panels), this fluorescence was intranuclear and did not accumulate along the nuclear periphery even at low levels of expression (see timecourse of cells treated with galactose, Extended Data Fig. 1a). We interpret these data in a model where Atg39 can cross the pore membrane, but is then liberated from the INM likely through an INM Associated Degradation (INMAD)-type mechanism^39–42^. Such a model predicts the existence of a degron sequence in Atg39, likely in its N-terminus. Consistent with this idea, deletion of the N-terminus of Atg39 resulted in the accumulation of a reconstituted nuclear rim fluorescence when GFP^1–10^-atg39-(139-398)(Fig. 1a) was expressed with the nucleoplasmic reporter, even after several hours of growth in galactose and thus high levels of expression (Fig. 1e,f, Extended Data Fig. 1b), whereas deletion of the C-terminal lumenal (L) domain comprising amino acids 169-398 (GFP^1–10^-atg39-ΔL)(Fig. 1a) mirrored the full-length protein, albeit with a more visible pool at the nuclear periphery at lower levels (Fig. 1g,h, Extended Data Fig. 1c). Thus, there are sequence elements in the N-terminus of Atg39 that might trigger its removal from the INM. Taken together, the data are most consistent with a model in which Atg39 localizes at the ONM and may in fact be restricted from accumulating at the INM.

### The Atg39 lumenal domain is required for ONM targeting and bleb formation

A model in which Atg39 acts from the outside of the nucleus – in predicts physical interactions that connect Atg39 to the INM through the NE lumen/perinuclear space. To explore this, we generated a deletion series of Atg39 to systematically evaluate the sequence elements that conferred ONM localization; we also evaluated the ability of these constructs to induce NE blebs. As a translumenal interaction is predicted to confer ONM accumulation, we first generated C-terminal deletions that sequentially removed lumenal regions that secondary structure prediction suggested were alpha (α) helical in nature (Fig. 2a). Deletion of the terminal two alpha helical segments (GFP-atg39-(1-333)) did not impact NE targeting nor the number of NE blebs observed in each cell (Fig. 2b,e). In contrast, the removal of α2 (GFP-atg39-(1-312)) led to a marked decrease in the number of NE blebs/cell with no obvious impact on NE targeting (Fig. 2b,e). These data suggest that the formation of NE blebs requires a discrete sequence motif in the Atg39 lumenal domain. They further imply that the NE remodeling observed is not simply an artifact of overexpression but may in fact represent a membrane deforming activity present in the Atg39 lumenal domain.

**Fig. 2.**
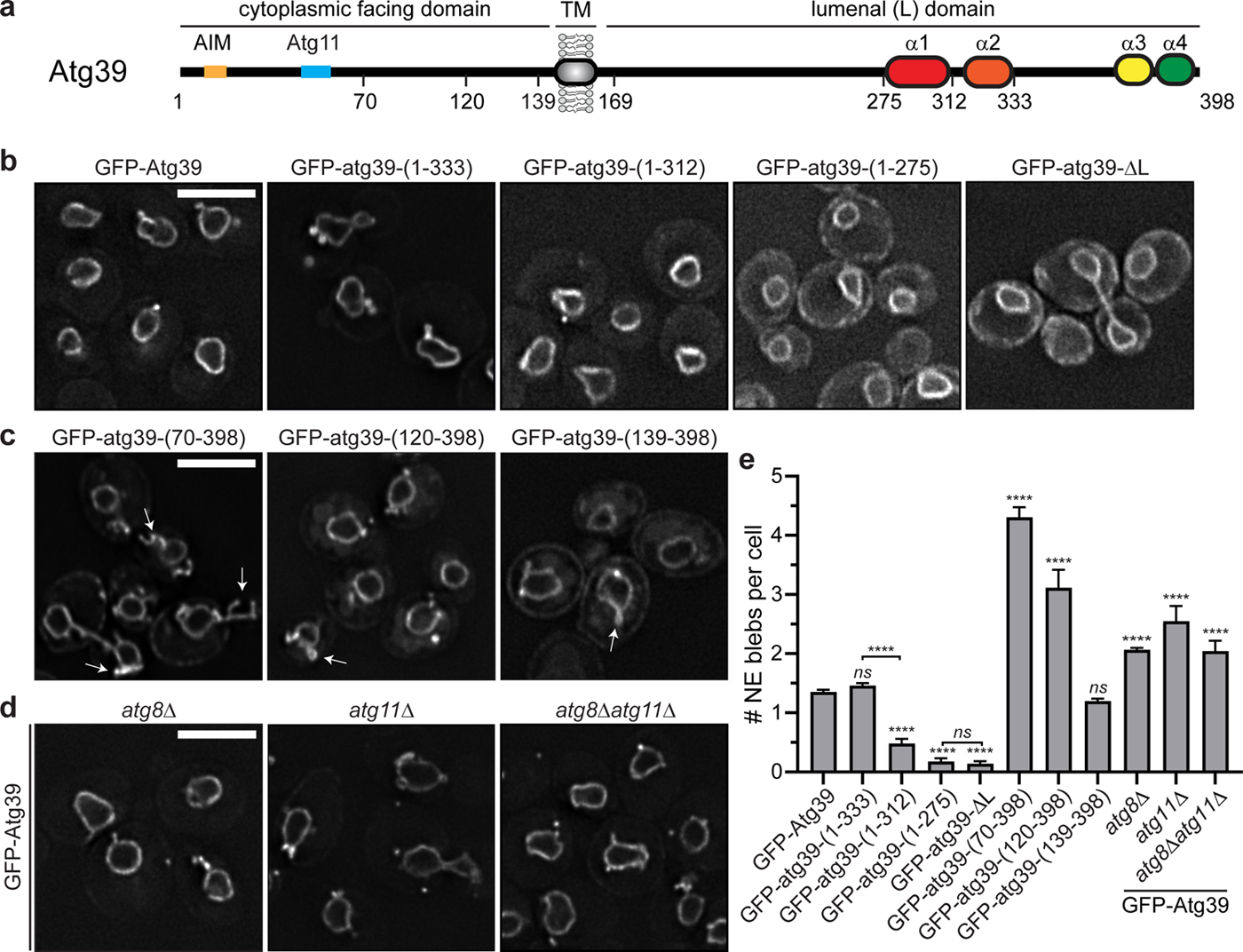
The Atg39 lumenal domain is required for NE targeting and bleb formation. **a**, Schematic of Atg39 with AIM (Atg8 interacting motif, light orange), Atg11-binding region (blue), predicted transmembrane (TM) helix (grey) and predicted alpha (α) helices 1-4 (colored ovals). Numbers are amino acids. **b**, Deconvolved fluorescence micrographs of the indicated GFP fusion constructs. Numbers indicate amino acid position from **a**. Scale bar is 5 µm. **c**, As in **b**. **d**, Deconvolved fluorescence micrographs of GFP-Atg39 in the indicated strains. Scale bar is 5 µm. **e**, Bar chart plotting the quantification of the number of NE blebs per cell in the indicated strains from micrographs in **b-d**. At least 50 cells of each genotype were quantified from three independent experiments. Average and SD are shown. Statistics are from one-way ANOVA where *ns* is *p* > 0.05, and **** *p* ≤ 0.0001.

Removal of α1 abrogated NE accumulation as both GFP-atg39-(1-275) and GFP-atg39-ΔL no longer exclusively accumulated at the NE and were found throughout the cortical ER as well (Fig. 2b). Thus, these data support a model in which the lumenal domain of Atg39 has sequence elements required for both NE accumulation (α1) and NE remodeling (α2). In analogy to KASH-proteins that accumulate at the ONM by binding to INM SUN proteins^43^, we propose that Atg39 accumulates at the ONM by forming a translumenal bridge through direct or indirect interactions with the INM.

NE targeting and remodeling activities appear to be largely restricted to the lumenal domain as N-terminal deletions of Atg39 did not appreciably impact the accumulation of GFP-atg39-(70-398), GFP-atg39-(120-398) or GFP-atg39-(139-398) at the NE, although we did observe more elaborate NE-blebs (Fig. 2c, arrows and Fig. 2e) in these cells that could be correlated to higher levels of these constructs (Extended Data Fig. 2a). This was intriguing as it suggested that interactions with the autophagy machinery, which are mediated through the Atg8 and Atg11-binding motifs^29^ (Fig. 2a) did not contribute to the formation of the NE-blebs. Consistent with this conclusion, similar numbers of GFP-Atg39 containing NE-blebs were observed in *atg8Δ* and *atg11Δ* strains compared to wild type cells (Fig. 2d,e) with GFP-Atg39 expressed at similar levels in these strains (Extended data Fig. 2b). Thus, the Atg39 lumenal domain is necessary for NE targeting and remodeling, which can occur independently of engagement with the core autophagy machinery.

### The Atg39 lumenal domain is required for nucleophagy

The data support a model in which Atg39 can accumulate and mediate NE remodeling from the ONM by virtue of sequence motifs in its lumenal domain. To evaluate the importance of these motifs in the nucleophagic clearance of INM proteins, we tested whether they were required for the autophagic degradation of the established integral INM Atg39-cargo, Heh1 (also called Src1)^29, 32^ under conditions of nitrogen starvation.

We tested the degradation of Heh1-GFP using a standard autophagy assay that relies on the visualization of a stable fragment of GFP (GFP’) by Western blot, which is liberated from Heh1-GFP by vacuolar proteases. Consistent with published data^29^, we observed a ∼65% degradation of the total pool of Heh1-GFP in cells grown in medium lacking nitrogen, which was mitigated in the *atg39Δ* strain (Fig. 3a,b). This effect was specific for Heh1-GFP as the relative degradation of Heh1’s paralogue, Heh2-GFP, and the nucleoporin Nup82-GFP, were not significantly impacted by *ATG39* deletion (Fig. 3c,d).

**Fig. 3.**
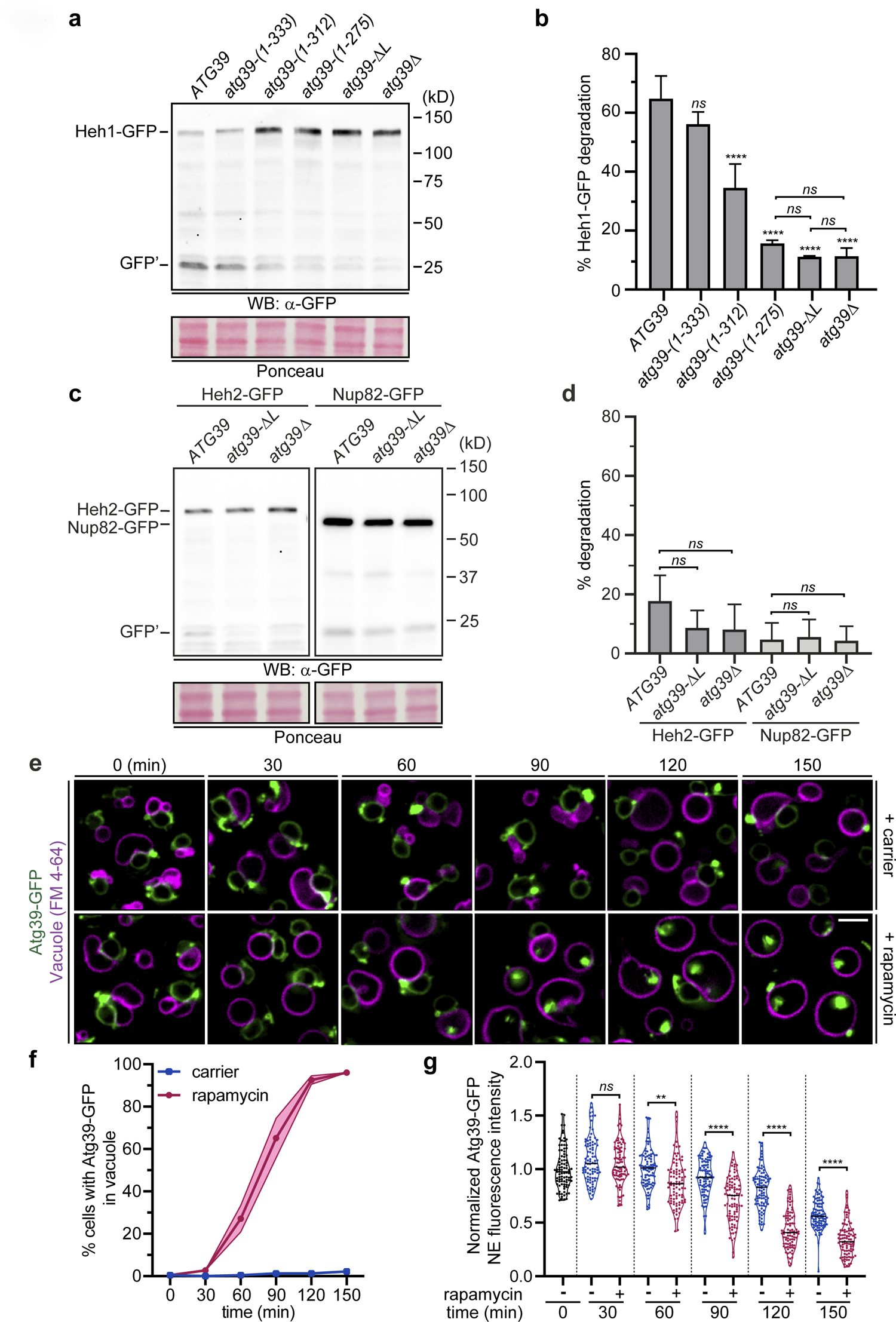
Atg39-containing NE-blebs are delivered to vacuoles by autophagy. **a**, The autophagic degradation of Heh1-GFP requires the Atg39 lumenal domain. Western blot of proteins (anti-GFP.1) from whole cell extracts derived from cells expressing Heh1-GFP in strains with the indicated *atg39* alleles treated with rapamycin. GFP’ is a stable fragment of GFP in vacuoles. Position of molecular weight standards (in kD) at right. To assess relative protein loading, a portion of the blot is shown stained with Ponceau. **b**, Plot of mean and SD of the percentage degradation of Heh1-GFP from three independent experiments as in **a**, Statistics from one-way ANOVA test and Tukey’s correction where *ns* is *p* > 0.05, and **** *p* ≤ 0.0001. **c**, As in **a** but examining autophagic degradation of Heh2-GFP and Nup82-GFP. **d**, As in **b** but plotting the percentage degradation of Heh2-GFP and Nup82-GFP. **e**, Deconvolved fluorescence micrographs of a timecourse (30 min intervals) of rapamycin or carrier-alone treated (DMSO) cells expressing Atg39-GFP (green) in a *pep4Δ* strain; vacuoles are stained with FM 4-64 (magenta). Merged fluorescent images shown. Scale bar is 3 µm. **f**, Line plot of percentage of cells where Atg39-GFP is visible in vacuoles after treatment with rapamycin (circles and magenta line) or carrier (squares and blue line) over the timepoints indicated. 75 cells were evaluated each from three independent replicates. SD from the mean percentage is indicated by the shaded area. **g**, Violin plot of the fluorescence intensity of Atg39-GFP along the nuclear periphery normalized to background fluorescence in the presence of rapamycin (+) or carrier (-) at the indicated timepoints. 30 cells each from three independent replicates were evaluated, each replicate was normalized to the mean NE fluorescence at 0 min. Solid line denotes the median, width of the violin plot denotes relative frequency of data points. Statistics are from Student’s t-test with Welches correction where *ns* is *p* > 0.05, ** is *p* ≤ 0.01 and **** is *p* ≤ 0.0001.

To test the role of the Atg39 lumenal domain motifs in nucleophagy, we next examined Heh1-GFP degradation in strains expressing lumenal truncations from the endogenous *ATG39* chromosomal locus (i.e. under the control of the *ATG39* promoter). Consistent with its importance for executing nucleophagy, the sequential trimming of the C-terminal lumenal domain resulted in a reduction of Heh1-GFP degradation that reflected the NE targeting and remodeling analysis (Fig. 2). For example, consistent with the dispensability of the terminal two α-helical segments for NE targeting and remodeling (Fig. 2b,e), the *atg39-(1-333)* allele fully complemented the degradation of Heh1-GFP as the wild type *ATG39* gene (Fig. 3a,b). In contrast, the sequential removal of the α2 (*atg39-1-312*) and α1 (*atg39-1-275*) coding segments, required for NE remodeling and NE targeting, respectively, resulted in a progressive loss of the ability of these alleles to contribute to Heh1-GFP degradation (Fig. 3a,b). Thus, the lumenal sequence motifs that are required for Atg39 NE targeting and blebbing are also needed to effectively execute nucleophagy under conditions of nitrogen starvation.

### Atg39-containing NE-blebs are delivered to the vacuole by autophagy

Because of the obvious overlap between the requirement of the lumenal motifs for NE targeting/remodeling and nucleophagy, we considered the possibility that the NE-blebs formed upon Atg39 overexpression may in fact represent a morphologically relevant intermediate in nucleophagy, albeit exaggerated due to its high levels. Were this to be the case, several criteria needed to be met. First, if the NE-blebs were an intermediate in nucleophagy, they would need to be delivered to the vacuole in a mechanism requiring core autophagy genes. Second, the expectation would be that the NE-blebs would contain cargo specific for Atg39-mediated nucleophagy. Third, the NE ultrastructure driven by Atg39 would need to reflect characteristics of protein-mediated membrane remodeling as opposed to simply membrane expansion or the formation of membrane stacks or lamellae that are common artifacts of the overexpression of NE and ER membrane proteins^44–52^.

To address these criteria, we first tested whether the NE-blebs were delivered to vacuoles. To induce autophagy, we treated Atg39-GFP expressing cells with rapamycin while arresting Atg39-GFP production by inhibiting the *GAL1* promoter with the addition of glucose to the medium; delivery of Atg39-GFP to vacuoles (labeled with the FM 4-64 dye) was monitored by fluorescence microscopy at 30 min intervals (Fig. 3e). To ensure that Atg39-GFP could be visualized in vacuoles, these experiments were performed in a strain lacking Pep4, a vacuolar protease required for activation of vacuolar hydrolases^53^. As shown in Fig. 3e, we observed internal vacuolar GFP fluorescence in ∼25 % of rapamycin-treated cells beginning at the 60 min time point, which progressively increased to ∼95 % by the end of the experiment (150 min; Fig. 3f). Consistent with the idea that Atg39-GFP is progressively being cleared by autophagy, we observed a coincident reduction in Atg39-GFP fluorescence at the nuclear periphery (Fig. 3g).

Further, we did not observe GFP fluorescence in the vacuole in carrier-alone (DMSO) treated samples despite some reduction to its levels at the nuclear periphery (Fig. 3e,f,g). We interpret the latter to reflect that Atg39-GFP production is inhibited upon addition of glucose to the medium and there is a dilution of Atg39-GFP signal through the ∼1.5 cell divisions that occur during the 150 min time course. Therefore, overexpressed Atg39-GFP can be delivered to the vacuole under conditions of autophagy induction by rapamycin treatment.

To further confirm that overexpressed Atg39-GFP was degraded by autophagy, we also observed the production of GFP’ after treating Atg39-GFP expressing cells with rapamycin (Extended Data Fig. 2c). Consistent with the conclusion that this GFP fragment was the product of a vacuolar protease, it was not observed in a *pep4Δ* strain (Extended Data Fig. 2c). Importantly, this autophagy-dependent degradation of Atg39-GFP also required the Atg39 lumenal domain as GFP’ was not visible upon treatment of atg39-ΔL-GFP expressing cells with rapamycin (Extended Data Fig. 2d). Lastly, we confirmed that Atg39-GFP is targeted by autophagy by testing the production of GFP’ in both *atg8Δ* and *atg11Δ* strains. Deletion of *ATG8,* and to a lesser extent *ATG11,* abolished its production (Extended Data Fig. 2e). We conclude that Atg39-GFP containing blebs can be delivered and degraded in the vacuole through an autophagy-dependent mechanism.

### Atg39-derived NE-blebs specifically capture the INM

The second criteria that would provide confidence that the NE-blebs are a potentially physiological intermediate in nucleophagy would be the selective incorporation of Atg39 cargo into the blebs. To test this, we monitored the distribution of several integral components of the NE (Fig. 4a) expressed at endogenous levels including NPCs (Nup85), spindle pole bodies (Spc42) and integral INM proteins Heh1 and Heh2, in the context of mCherry-Atg39 expression. Of these, only Heh1 has been shown to be targeted by Atg39-dependent nucleophagy^29, 32^, whereas Nup85 can be degraded by NPC-phagy^17^. And indeed, we did not observe any appreciable accumulation of NPCs within the NE-blebs, nor components of the SPB (Fig. 4b, right “GFP” panels, 4c). In striking contrast, Heh1-GFP co-localized with mCherry-Atg39 within at least 50% of the blebs (Fig. 4b, arrowheads, Fig. 4c). As Heh1-GFP is expressed at low levels^54^ often below our threshold for detection even at the NE, we suspected that these numbers were an underestimate. In fact, Heh1-GFP is produced from one of two alternatively spliced forms of the *HEH1-GFP* transcript^55^. By examining the localization of a truncation made before the splice site, a more abundant INM-localized heh1-ΔL-GFP (Fig. 4a) can be visualized in virtually all Atg39-containing NE-blebs (Fig. 4b,c). Further, the brighter heh1-ΔL-GFP allowed the continual monitoring of heh1-ΔL-GFP sequestration into the blebs by timelapse microscopy, which occurs concomitantly with bleb formation (Extended Data Fig. 3, arrowheads).

**Fig. 4.**
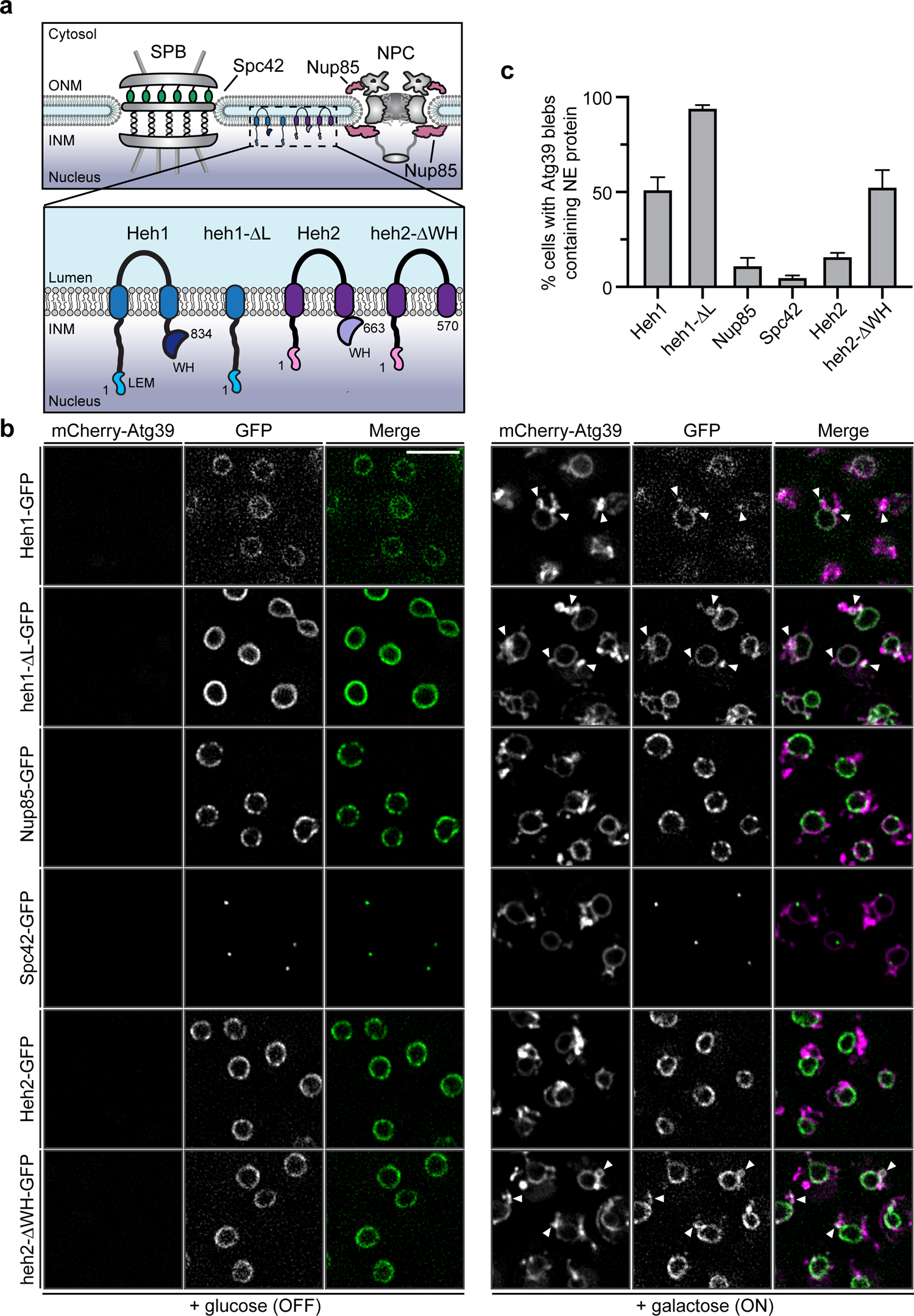
INM is specifically captured in Atg39-containing NE blebs. **a**, Cartoon of protein and protein complexes at the NE including a spindle pole body (SPB) and nuclear pore complex (NPC). Boxed region is magnified at bottom to show integral INM proteins and truncation mutants. LEM is LAP2/emerin/MAN1, WH is winged helix (WH). Numbers are amino acids. **b**, Deconvolved fluorescent micrographs of the indicated GFP fusion proteins under conditions where mCherry-Atg39 expression is repressed (+ glucose, OFF; left panels) or induced (+ galactose, ON; right panels). Merged image of mCherry (magenta) and GFP (green) is also shown. Arrowheads point to NE blebs containing mCherry-Atg39 and INM proteins. Scale bar is 5 µm. **c**, Quantification of percentage of cells with Atg39 blebs colocalized with indicated NE proteins. Error bars are SD from three independent replicates of 100 cells per replicate.

Surprisingly, the paralogue of Heh1, Heh2, was not captured with Atg39 in the blebs (Fig. 4b,c) suggesting that there may be some selectivity for subdomains of the INM. To explain this observation, we recently uncovered that a substantial fraction of Heh2 is bound to NPCs^56^, which might prevent Heh2’s capture into the NE blebs. To test this idea, we examined the localization of a truncation of Heh2 (heh2-ΔWH; Fig. 4a) that retains INM-targeting information but lacks the ability to bind NPCs, which is conferred by a C-terminal winged-helix (WH) domain^56^. Consistent with the conclusion that binding to NPCs prevents capture of Heh2, heh2-ΔWH-GFP was found in 50% of the visible Atg39-containing NE-blebs (Fig. 4b,c). Thus, taken together, Atg39-dependent NE-blebs can selectively capture the INM over other elements of the NE suggesting that the overexpression of Atg39 might be recapitulating key early steps in an Atg39-dependent nucleophagy pathway.

### NE-blebs contain a network of INM-derived vesicles in the NE lumen

Lastly, to evaluate whether Atg39 overexpression leads to changes in NE morphology that might illuminate early steps in nucleophagy, we turned to CLEM and tomography. We first examined cells prepared from cultures expressing high levels (Extended Data Fig. 4a) of Atg39-GFP (Fig. 5a, Extended Data Fig. 4b, Supplementary Video 1).

**Fig. 5.**
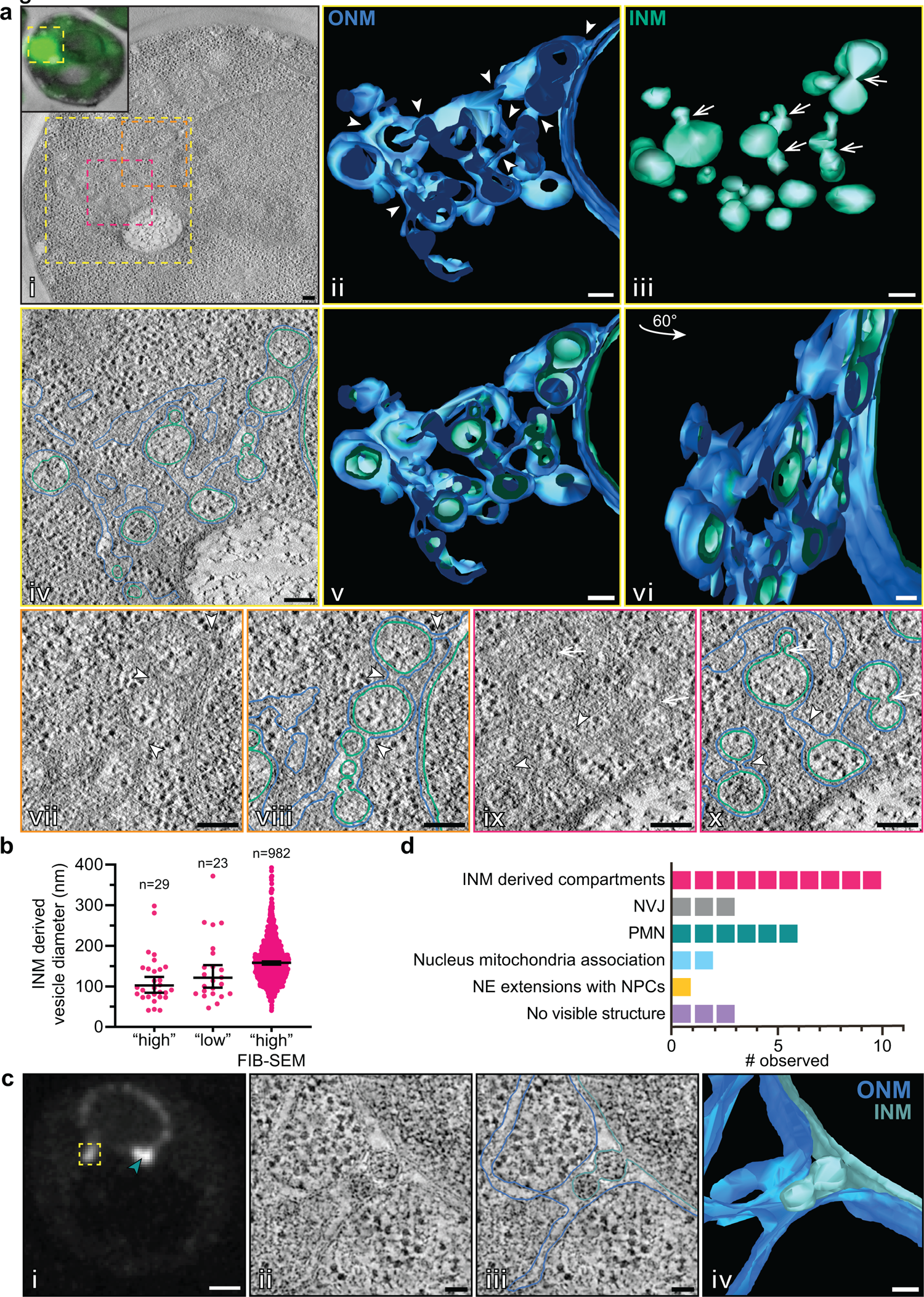
CLEM of Atg39-GFP reveals expanded ONM with INM-derived vesicles in the NE lumen. **a**, (i) Virtual slice of an electron tomogram of cell with “high” expression of Atg39-GFP (fluorescence image overlaying EM in inset). Boxes represent magnified regions shown in the following panels with corresponding colors. (ii) 3D model of continuous ONM with arrowheads pointing to continuities between substructures. (iii) 3D model of likely INM-derived vesicles within the ONM compartment with arrows pointing to constrictions. (iv) Example virtual slice of tomogram with ONM and INM traced in blue and teal, respectively. (v, vi) Two views of 3D model where ONM and INM are segmented. (vii, viii and ix, x) Further magnifications of single tomographic slices (with and without annotation) as defined by surrounding box color with key in (i). Arrowheads point to continuity of ONM and arrows to INM constrictions. Scale bars are 200 nm. **b**, Quantification of the diameter of INM-derived compartments from cells with “high expression (4 cells), “low” expression (10 cells) of Atg39 from electron tomography or “high” expression of Atg39 from FIB-SEM (52 cells). Total number of INM vesicle diameters measured is indicated in the figure. Median and 95% confidence interval are shown. **c**, CLEM of “low” expressed GFP-Atg39. (i) Fluorescence micrograph of GFP-Atg39 with boxed region ultrastructure shown in (ii). Arrowhead points to a region of PMN. (ii) Virtual slice from electron tomogram acquired at of region indicated by the box in panel (i). (iii) Annotation of virtual slice from panel (ii) with the ONM in blue, the INM in teal. (iv) 3D surface rendering of annotated structures in panel (iii). Scale bars: (i), 2 µm; (ii– iv), 50 nm. **d**, Quantification of indicated morphologies/substructures observed from 26 total GFP-Atg39 NE foci.

Correlation of regions of Atg39-GFP fluorescence (Fig. 5a, Extended Data Fig. 4b, insets) with their position in electron micrographs revealed an impressive proliferation of membranes. These membranes were derived from the NE as direct continuity could be observed with the ONM in single tomographic slices and in 3D reconstructions; ONM ultimately enclosed the entire structure (Fig. 5a, ii, iv-x, arrowheads point to ONM continuity). Most strikingly, captured within the expanded ONM were additional bilayers that are traced in teal (Fig 5a, iv, viii, x), which were most logically derived from the INM. As these membranes were circular within single tomographic slices, we speculated that they were vesicles. Consistent with this, a 3D model of these segmented membranes revealed that they were spherical and similarly sized with a median diameter of ∼115 nm (Fig. 5a, iii, 5b). Interestingly, a subset of vesicles were connected by membrane constrictions as if undergoing fission (Figure 5a, iii, ix and x, arrows).

The vesicle diameter measurements were likely an underestimate as it was not possible to capture a large number of entire vesicle volumes within the relatively thin ∼200 nm thick tomograms. We therefore also performed FIB-SEM on Atg39-GFP expressing cells. This approach allowed the visualization of 52 whole cell volumes (Extended Data Fig. 5a) revealing cells with expansive networks of NE blebs that emanated from multiple sites on single nuclei (Extended Data Fig. 5b,c, Supplementary Video 2). In these 3D images, we measured the diameter of 982 INM derived vesicles, which had a median diameter of ∼164 nm (Fig. 5b). Interestingly, we also observed lipid droplets associated with virtually all of the NE blebs (Extended Data Fig. 5b,c). Thus, the overexpression of Atg39 leads to the generation of a network of likely INM-derived vesicles within the NE lumen alongside an expansion of the ONM with associated lipid droplets. While there are certainly many examples of overexpressed membrane proteins driving changes to membrane morphology, the observed Atg39-dependent morphology is most analogous to those observed in NE egress pathways^57^.

To gain more insight into the biogenesis of the Atg39-induced compartments, we next performed CLEM on cells expressing lower levels of a GFP-Atg39 fusion (Extended data Fig. 4a). We examined the ultrastructure at NE sites of local GFP-Atg39 accumulation and where emerging blebs were visible by fluorescence microscopy (Fig. 5c, Extended data Fig. 4c-f). Ultimately, we examined 26 individual Atg39 focal NE accumulations within 23 cells. Of these, 3 could not be attributed to any obvious morphology with 2 being localized at sites where mitochondria were adjacent to the NE (Fig. 5d, Extended data Fig. 4c). In 9 cells, we could unambiguously correlate the fluorescence to either NVJs (3 out of 9) or regions of PMN (6 out of 9)(Fig. 5d, Extended Data Fig. 4d), which may be consistent with recent work supporting a role for Atg39 in PMN^33^. And, in a single cell we observed likely extensions to the NE that contained NPCs (Fig. 5d, Extended data Fig. 4e).

The most prevalent morphology (10 out of 26; Fig. 5d) correlated with GFP-Atg39 is presented in Fig. 5c and Extended Data Fig. 4f. As shown in Fig. 5c, we observed direct continuity between the INM and vesicles within the NE lumen with a ∼25 nm constriction or bud neck at the INM. As in the high level expression scenario, these vesicles were again similarly sized (median diameter of 139 nm, Fig. 5b) and were sometimes found in a series where membrane connections could be visualized, segmented and 3D modeled (Fig. 5c panel, iii, iv, Supplementary Video 3). Simultaneously, we observed a clear expansion of the ONM, presumably necessary to accommodate the presence of the extra volume occupied by the vesicles in the NE lumen. Thus, we interpret these structures as precursors to the more elaborate compartments observed upon high level expression of Atg39. These data suggest that Atg39 may have the ability to coordinate membrane remodeling between the INM and ONM and capture components of the INM into vesicles in the NE lumen.

## Discussion

Atg39 has recently emerged as a key player in the autophagic degradation of nuclear and INM components^17, 29–32, 58^, however, how nuclear cargo is selectively packaged and delivered to the cytosolic autophagosome, through the double membraned NE, remained open questions. Here, we provide a framework for answering these questions by proposing an outside-in model of nucleophagy that depends on Atg39 acting from its position at the ONM (see model, Extended Data Fig. 5d). There, Atg39 can engage with the cytosolic autophagy machinery through its cytosolically-exposed N-terminal domain while connecting to the INM through its lumenal domain. Evidence in support of this model includes the demonstrated importance of the Atg39 lumenal domain not only for NE targeting and NE remodeling (Fig. 2) but also for the nucleophagic degradation of a model Atg39 INM cargo, Heh1-GFP (Fig. 3). Such a mechanism may prove relevant to other forms of nuclear autophagy as well. For example, PMN requires the generation of an ONM-vacuole contact site by the pairing of Nvj1 and Vac8^59^: the close apposition of the INM and ONM at these sites is thought to be mediated by the Nvj1 lumenal domain^60^. Interestingly, however, Nvj1 overexpression does not lead to any obvious membrane deformation^59^. Therefore, recent evidence supporting a role for Atg39 in PMN^33^ might suggest a model in which Atg39 contributes a membrane remodeling activity capable of co-evaginating the INM and ONM, which is a pre-requisite for both nucleophagy mechanisms.

Perhaps the most exciting element of this model is the proposal that the INM and associated nuclear content is captured within NE lumenal vesicles derived from the INM (Fig. 5; Extended Data Fig. 4,5). Although we acknowledge that these structures are observed upon Atg39 overexpression, we argue that they are likely bona fide early intermediates in a physiological nucleophagy mechanism. Our confidence with this conclusion is based on several data. First, there is the remarkable ability of the Atg39-driven NE-blebs to selectively capture established INM cargo over other components of the NE (Fig. 4). Second, the NE-blebs can be targeted by autophagy provided the signal, yet to be defined, is supplied (Fig. 3e,f). These data also suggest that the interaction between Atg39 and the autophagy apparatus may be regulated in some way, a common theme with selective autophagy cargo adaptors^61, 62^. Third, the overexpression of Atg39 lacking lumenal elements, in particular the predicted α helix 2, does not drive analogous morphologies (Fig. 2), further fortifying that they are a specific consequence of Atg39 and not simply an accumulation of an overexpressed integral NE protein.

The enclosure of INM within lumenal vesicles is also attractive because it provides a harmonious mechanism for how the INM could be selectively removed without impacting NE integrity. The observed ∼25 nm membrane necks where the vesicle membranes connect with the INM are strongly suggestive of a protein-scaffold that would maintain their stability and ultimately drive membrane scission. This observation, in addition to the chain-like concatenation of the INM-derived vesicles, strongly resembles morphologies associated with ESCRT function^63, 64^. The hypothesis that ESCRTs may ultimately be involved in a scission event that would liberate the INM-derived vesicle into the lumen is attractive not only because of the many links between autophagy and ESCRTs^44, 65–69^, but also because Heh1, the only well-established INM protein cargo of Atg39^29^, directly engages with ESCRTs in pathways that ensure NE integrity^70, 71^.

Lastly, the removal of INM contents through the proposed mechanism evokes comparisons to the nuclear to cytosolic translocation of so-called “Mega” RNPs in some *Drosophila* neurons that proceeds through a vesicular intermediate in the NE lumen^72^. Thus, we anticipate that the removal of intranuclear contents through the NE lumen will prove to be a generalizable principle of protein, and perhaps RNA, quality control, that will be relevant beyond the yeast system and has in fact already been hypothesized^73^. Ultimately understanding whether this is the case will require the identification of a mammalian functional homologue to Atg39, which so far remains elusive but is the focus of active investigation.

## Acknowledgements

We thank members of the Melia and LusKing labs for discussion and feedback. We are indebted to Morven Graham, Xinran Liu for invaluable EM expertise. We thank M.P. Rout and S.L. Jaspersen for reagents. This work was funded by grants from the NIH (R21 AG058033 to C.P.L. and T.J.M.; R01 GM105672 to C.P.L.; F32 GM139285 to N.R.A.).

## Author Contributions

Conceptualization: S.C., P.J.M., D.J.T., T.J.M., C.P.L.; Methodology: S.C., P.J.M., D.J.T., N.R.A.; Investigation: S.C., P.J.M., D.J.T., N.R.A.; Validation: S.C., P.J.M., D.J.T., N.R.A.; Formal analysis; S.C., P.J.M., D.J.T., N.R.A.; Writing – original draft: S.C., P.J.M., D.J.T., T.J.M., C.P.L.; Writing – review & editing: All authors; Supervision: M.C.K., T.J.M., C.P.L.; Funding acquisition: T.J.M., C.P.L.

## Competing interests

The authors declare no competing interests.

**Supplementary Video 1-** Related to Fig. 5a. “High” expression of Atg39-GFP results in the formation of NE-blebs. Virtual slices from electron tomograph and 3D surface rendering from tracing throughout the tomography with the ONM (blue) and INM (teal) are shown. Scale bar is 100 nm.

**Supplementary Video 2** – Related to Extended Data Fig. 5c. Visualization of whole volumes of cells expressing Atg39-GFP by FIB-SEM with 3D surface rendering of membranes. The NE is shown in blue, lipid droplets in yellow, vacuole in grey, and mitochondria in dark pink. Scale bar is 200 nm.

**Supplementary Video 3** – Related to Fig. 5d. “Low” expression of GFP-Atg39 leads to formation of INM derived vesicles in the NE lumen. Virtual slices from an electron tomogram and 3D surface rendering from tracing throughout the tomography with the ONM in blue and INM in teal. Direct continuity between the INM and INM-derived vesicle necks are visible. Scale bar is 50 nm.

## Materials and Methods

### Yeast strain construction and culturing conditions

All strains used in this study are listed in Supplementary Table 1. Genomic integration of sequences encoding fluorescent reporter genes, replacement of endogenous gene promoters with G*AL1* promoter and gene deletions were generated using a PCR-based homologous recombination approach using the pFA6a plasmid series^74, 75^ (as templates. Yeast were cultured to mid-log phase in YP (1% Bacto-yeast extract (BD), 2% Bacto-peptone (BD), 0.025% adenine hemi-sulfate (Sigma)) supplemented with 2% raffinose (R; BD), 2% D-galactose (G; Alfa Aesar) 2% D-glucose (D; Sigma). To maintain selection of plasmids, cells were cultured in Synthetic Complete (SC) medium (Sunrise Science) that lacked the indicated amino acids. All experiments were performed at 30°C. Standard protocols for transformations, mating, sporulation, and dissection were followed^76^.

For induction of autophagy using rapamycin, rapamycin (in DMSO; Sigma-Aldrich) or an equivalent volume of DMSO (carrier) was added to mid-log phase cultures to a final concentration of 250 ng/mL. Samples collected at time points indicated in the figures were prepared for imaging or immunoblotting as described below.

To induce autophagy by nitrogen starvation, mid-log phase cells were pelleted at ∼375 *g*, washed twice with Synthetic Defined (SD) lacking nitrogen (SD-N) medium (0.17% Difco Yeast Nitrogen Base without amino acids and ammonium sulfate (BD), and 2% D), resuspended in SD-N, and returned to a shaking incubator at 30°C for the amount of time indicated in figure legends.

To assess the localization of Atg39-GFP and GFP-Atg39 under the control of the *GAL1* promoter, strains (SCCPL39, SCCPL40, SCCPL80, SCCPL82, SCCPL95, SCCPL131, DTCPL911, PMCPL87, PMCPL112, PMCPL113, PMCPL114, PMCPL115, PMCPL28, PMCPL29, PMCPL390, PMCPL392, PMCPL422, PMCPL424, and PMCPL471) were grown in YPR to mid-log phase. Expression was induced by the addition of 2% G and images were acquired at timepoints indicated in the figures.

To examine the subcellular localization of Atg39 in the split-GFP assay, strains containing Atg39 split-GFP fusions under the control of the *GAL1* promoter (PMCPL21, PMCPL34, PMCPL35, and PMCPL298) were transformed with plasmids containing split-GFP-mCherry reporters for the nucleoplasm (pSJ1321), ONM/ER (pSJ1568) or lumen (pSJ1602). Cells grown overnight in media lacking leucine were diluted into YPAR. Expression of Atg39 fusions was induced by the addition of 2% G for 4 h or as otherwise indicated in the figure legends.

To visualize Atg39-GFP within vacuoles upon induction of autophagy, strain SCCPL22 was treated with 2% G to induce Atg39 overexpression. In order to visualize the vacuole membrane, 10 ml of culture was transferred to a foil wrapped tube and incubated with FM 4-64 (1 µM in DMSO, Molecular Probes) for 30 min at 30°C. Cells were then pelleted and resuspended in YPG. Atg39 expression was arrested after 4 h with the addition of 2% D and samples were split. One culture was treated with rapamycin (final concentration, 250 ng/ml, Sigma-Aldrich) to induce autophagy and the other with DMSO (carrier).

### Plasmid generation

All plasmids used in this study are listed in Supplementary Table 2.

To generate pPM1 (pFA6a-TRP1-GAL1-GFP^1–10^), the region encoding GFP^1–10^ was PCR amplified (KOD polymerase, EMD Millipore) from pSJ1256 using primers containing the *Pac*I and *Asc*I (New England BioLabs) restriction sites. The ampliconwas restriction digested with *Pac*I/*Asc*I (New England BioLabs), gel purified (Qiagen) and ligated (T4 ligase, Invitrogen) into gel purified pFA6a*-*TRP1-GAL1 digested with *Pac*I/*Asc*I (New England BioLabs). Successful subcloning was confirmed by sequencing.

To generate pFA6a-3xHA-mCherry-natMX6, the 3xHA epitope sequence was PCR-amplified with Q5 DNA polymerase using pFA6a-3xHA-his3MX6 (Longtine et al., 1998) as a template. The PCR product was assembled into pFA6a-GFP-his3MX6 (Longtine et al., 1998) digested with *Sal*I and *Pac*I (New England BioLabs) using the Gibson Assembly Master Mix (New England BioLabs).

To generate pSC8, the 5’ sequence of the native promoter and amino acids 1-443 of *HEH1* were amplified via PCR (KOD polymerase, EMD Millipore) from genomic DNA using primers encoding the restriction sites *BamH*I and *Hind*III. The subsequent fragment was gel purified (Qiagen) and ligated using T4 ligase (Invitrogen) into pRS415-GFP plasmid linearized with *BamH*1-HF and *Hind*III-HF. The C-terminal region of Heh1 was subsequently introduced by annealing oligos encoding *HEH1* amino acids 449-477 flanked with restriction sites *Hind*III and *Sal*I, followed by ligation using T7 ligase (Invitrogen). The entire fragment GFP-heh1(1-443)-HindIII-heh1(449-477) was excised by restriction digest with *BamH*I-HF and *Sal*I-HF. This purified fragment was ligated into *pRS405* digested with *BamH*I-HF and *Sal*I-HF. pSJ1602 (pRS315-NOP1pr-mCherry-SCS2TM-GFP11) was a gift from Sue Jaspersen (Addgene plasmid # 86417; http://n2t.net/addgene:86417; RRID:Addgene_86417). pSJ1321 (pRS315-NOP1pr-GFP11-mCherry-PUS1) was a gift from Sue Jaspersen (Addgene plasmid # 86413; http://n2t.net/addgene:86413; RRID:Addgene_86413). pSJ1568 (pRS315-NOP1pr-GFP11-mCherry-SCS2TM) was a gift from Sue Jaspersen (Addgene plasmid # 86416; http://n2t.net/addgene:86416; RRID:Addgene_86416). pSJ1256 (pFA6a-link-yGFP1-10-CaURA3MX) was a gift from Sue Jaspersen (Addgene plasmid # 86419; http://n2t.net/addgene:86419; RRID:Addgene_86419).

### Microscopy

For all live-cell imaging, mid-log phase cells were gently pelleted and washed with SC media contain 2% D and immediately imaged directly on cover glass. All images were acquired on an Applied Precision DeltaVision microscope (GE Healthcare Life Sciences) equipped with a 100x 1.4 NA oil immersion objective (Olympus), solid state illumination, a CoolSnapHQ^2^ CCD camera (Photometrics) or EMCCD (Photometrics). The microscope stage was maintained at 30°C within an environmental chamber.

### Image processing and analysis

All presented micrographs were deconvolved using an iterative algorithm in SoftWoRx (6.5.1; Applied Precision, GE Healthcare). Micrographs and immunoblots were analyzed in FIJI/imageJ^77^. Unprocessed images were used for quantification of fluorescence intensity.

Line profiles of fluorescence intensity were generated using the Plot Profile function in FIJI/ImageJ^77^. The minimum and maximum measured values from individual fluorescent channels were normalized to 0 and 1, respectively.

To quantify changes in Atg39-GFP fluorescence intensity at the NE, the integrated density of a region of interest at the nuclear periphery comprising 4-pixels was measured and normalized to mean background fluorescence.

### Secondary Structure prediction

The secondary structure of Atg39’s lumenal domain was predicted by threading the amino acid sequence of Atg39 through Jpred4^78^.

### Statistical methods

Graphs were generated using Prism (GraphPad 9.0). Statistical significance tests were used as indicated in figure legends. Significance values were calculated within Prism (GraphPad 9.0) and p-values are indicated on the graph or in figure legends as: *ns*, *p* > 0.05; * *p* ≤ 0.05; ** *p* ≤ 0.01; *** *p* ≤ 0.001; **** *p* ≤ 0.0001. Error bars are described in figure legends.

### Correlative light and electron microscopy

Correlative microscopy of resin-embedded cells was performed as previously described^79^. Expression of Atg39 fusion proteins from cells (DTCPL688 (Atg39-GFP) and PMCPL87 (GFP-Atg39)) cultured in YPAR, was induced by the addition of 2% G for 3 h. Cells were then collected by centrifugation for 2 min at 350 g. Yeast slurry was transferred to the 200 µm recess of an aluminum platelet (Engineering Office M. Wohlwend) and placed in an HPM100 (Leica Microsystems) for high pressure freezing. Samples were freeze-substituted in 0.1% uranyl acetate in acetone and embedded in Lowicryl HM20 (Polysciences) using the automated temperature control of an EM-AFS2 (Leica Microsystems) with manual agitation and solution exchange following the published protocol^80^. Resin was polymerized under UV light, and the resin-embedded cells were cut into 250 nm thick sections using an ultramicrotome (Leica Artos 3D) equipped with a diamond knife (Diatome) and collected on 200 mesh copper grids with carbon support (Ted Pella, Prod. # 01840).

Fluorescence and brightfield micrographs of resin-embedded sections were acquired as described above. Several Z-sections were acquired every 200 or 250 nm at each grid square of interest, and in-focus planes were selected for CLEM alignment and presentation in figures.

Grids selected for tomography were post-stained with uranyl acetate and lead citrate. 15 nm protein A-coated gold beads (CMC UMC Utrecht) were adhered to the top and bottom surfaces of grids and used as fiducial markers for the alignment and reconstruction of the tilt series. Single (Fig. 5a, Extended Data Fig. 4b-f) or double (Fig. 5c) axis tilt series were collected on a FEI TF20 electron microscope operating at 200 kV using a high-tilt tomography holder (Fichione Instruments; Model 2020) from approximately −65° to +65° with acquisition at 1° intervals. Images were acquired using SerialEM software^81^, at a 2 x 2 binned pixel size of 1.242 nm using a 4k x 4k Eagle CCD (FEI) camera with a 150 µm C2 aperture and a 100 µm objective aperture. Subsequent reconstruction and segmentation were completed in IMOD^82^ in an automated fashion^83^. For all virtual slices presented, a Gaussian filter in IMOD was applied to reduce noise.

Alignment of fluorescence and electron microscopy data was completed using the ec-CLEM Plugin^84^ in the ICY imaging suite^85^. Low magnification EM was related to fluorescence and brightfield micrographs by selecting ∼6-8 points that corresponded to features visible in both images.

### FIB-SEM

For visualization of entire cellular volumes using FIB-SEM, unfixed yeast slurries loaded into 200 µm aluminum hats were frozen using a Leica HMP100 at 2000 psi. The frozen samples were then freeze substituted using a Leica Freeze AFS unit starting at −95°C using 0.1% uranyl acetate in acetone for 50 h to −600°C, then rinsed in 100% acetone and infiltrated over 24 h at −600°C with Lowicryl HM20 resin (Electron Microscopy Science). Samples were placed in gelatin capsules and UV hardened at −450°C for 48 h. The blocks were cured for a further few days before the resin block was trimmed to rough area of interest and the surface cleanly cut using a Leica UltraCut UC7. The entire block was carefully removed with a fine blade, and mounted on an aluminum stub using conductive carbon adhesive and silver paint (Electron Microscopy Sciences, Hatfield, PA, U.S.A,), then sputtered coated with approximately 20 nm Pt/Pd (80/20) using Cressington HR equipment (Ted Pella,Inc. Redding CA) to reduce charging effects.

A dual beam FIB-SEM (Zeiss CrossBeam 550) using a Gallium ion source was used to mill and SE2 secondary electron detector was used to image the samples. SmartSEM (Zeisss, Jenna Germany) was used to set up initial parameters and to find the regions of interest (ROI) by SEM images at 10 kV, 50 µm width and 30 µm height. The actual depth was 30 µm with 7 nm per pixel and 7 nm per slice. A Platinum-protective layer was deposited at the ROI with the FIB (30 kV, 3 nA) to protect the structure and reduce charging. Milling and highlight was done at 30 kV and 50 pA, with a carbon deposit 30 kV 3 nA. A course trench milled 30 kv 30 nA followed by fine milling at 30 kV 3 nA and for final acquisition a cuboid the area of interest was milled at 30 kV and 300 pA. After milling each slice an image was taken by detecting backscattered electrons of a primary electron beam (acceleration voltage of 1.5 kV, imaging current of 2nA, and aperture diameter of 100 µm) with a pixel dwell time of 3 µs. Atlas5 (Zeiss) was used for preliminary SEM stack alignment and FIB/SEM image stacks were saved as TIFF and MRC files. The images were imported into Dragonfly software (ORS, Montreal Canada) for further alignment and segmentation. The total volume imaged was 19.25 µm x 35.8 µm x 8.9 µm. Segmentation of structures was done in IMOD^82^, approximately every 35 nm.

### Immunoblotting

For immunoblotting, cells were harvested as previously described^86^. Briefly, ∼1.5×10^8^ cells were treated with 10% TCA for 1 h on ice and centrifuged at 15,000 *g* for 10 min at 4°C. The pellet was washed with ice-cold acetone, homogenized by sonication (Bioruptor UCD-200) and pelleted by centrifugation. After two cycles of washing and sonication, the pellet was vacuum-dried for 15 min. The dried cell pellet was then mechanically disrupted with 100 µl glass beads (Sigma) and 100 µl urea cracking buffer (50 mM Tris-HCl pH 7.5, 8 M Urea, 2% SDS, and 1 mM EDTA), followed by addition of 100 µl protein sample buffer (Tris-HCl pH 6.8, 7 M urea, 10% SDS, 24% glycerol, bromophenol blue, and 10% β-mercaptoethanol).

To assess autophagy through GFP-fallout experiments, ∼1.5×10^8^ cells were harvested and suspended in 0.2 M NaOH containing 0.1 M DTT, incubated on ice for 10 min, followed by the addition of 10% TCA with incubation on ice for 15 min. After centrifugation at 15,000 g for 5 min at 4°C, the pellet was washed with acetone, vacuum dried, and then resuspended in SDS sample buffer (0.1M Tris-HCl pH 7.5, 2% SDS, 10% glycerol, 20 mM DTT) for 10 min at 65°C to dissolve the pellet, followed by 95°C for 3 min.

Proteins from whole cell extracts were resolved on 4-20% SDS-polyacrylamide gels (BioRad, #4561096), followed by transfer of the proteins to 0.2 µm nitrocellulose membranes (Bio-Rad). The membranes were blocked in 5% non-fat milk in TBST for 1 h and immunoblotted with antibodies against GFP (anti-GFP.1, 4°C overnight, Takara Bio Clontech, 632381 or anti-GFP.2, 1 h room temperature, anti-GFP gift from M. Rout, as indicated in figure legends). Blots were incubated with HRP-conjugated secondary antibodies (1 h room temperature; Sigma) and visualized by ECL (Thermo Fisher Scientific) using a VersaDoc Imaging System (Bio-Rad). Relative protein loading was visualized using Ponceau S Solution (Sigma).

### Quantification of autophagic turnover

To calculate the relative percent degradation of GFP-fusion proteins, ROIs were drawn around immunoblot bands corresponding to cleaved GFP′ bands and GFP fusion proteins in Fiji/ImageJ^77^ and the total fluorescence intensity was measured. Measured values for GFP’ were divided by the sum of GFP′ and related GFP-fused full-length protein intensities.

---

**Extended Data Fig. 1.**
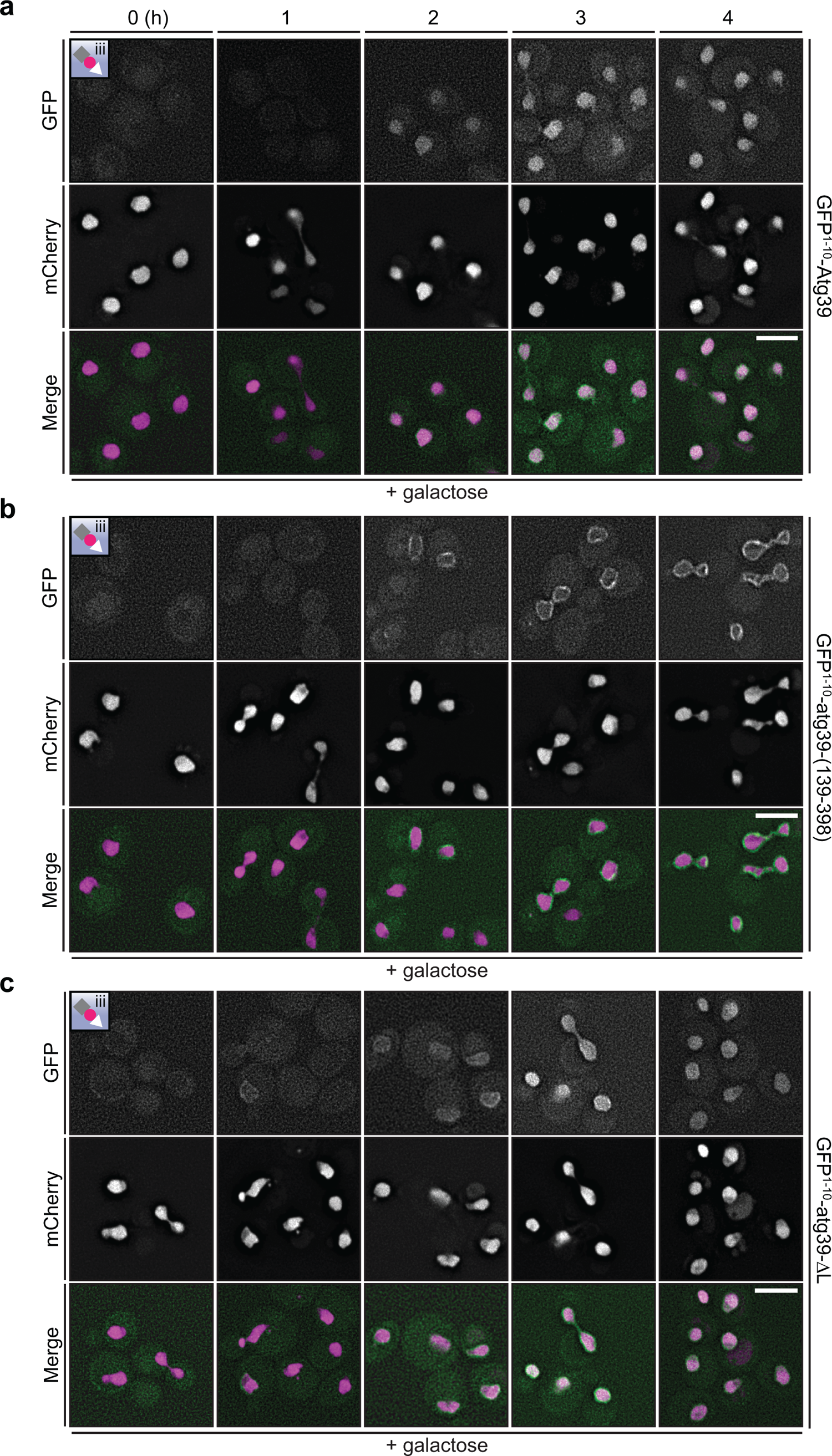
The split-GFP reconstitution of intranuclear GFP fluorescence depends on the N-terminus of Atg39. **a-c**, Deconvolved fluorescence micrographs of the indicated GFP^1–10^-constructs co-expressed with the nucleoplasmic split-GFP reporter (see inset and Fig. 1a). GFP^1–10^-constructs are expressed behind a *GAL1* promoter that is induced by growth in galactose for the indicated times (in hours). GFP (green), mCherry (magenta) and merged images shown. Scale bars are 5 µm.

**Extended Data Fig. 2.**
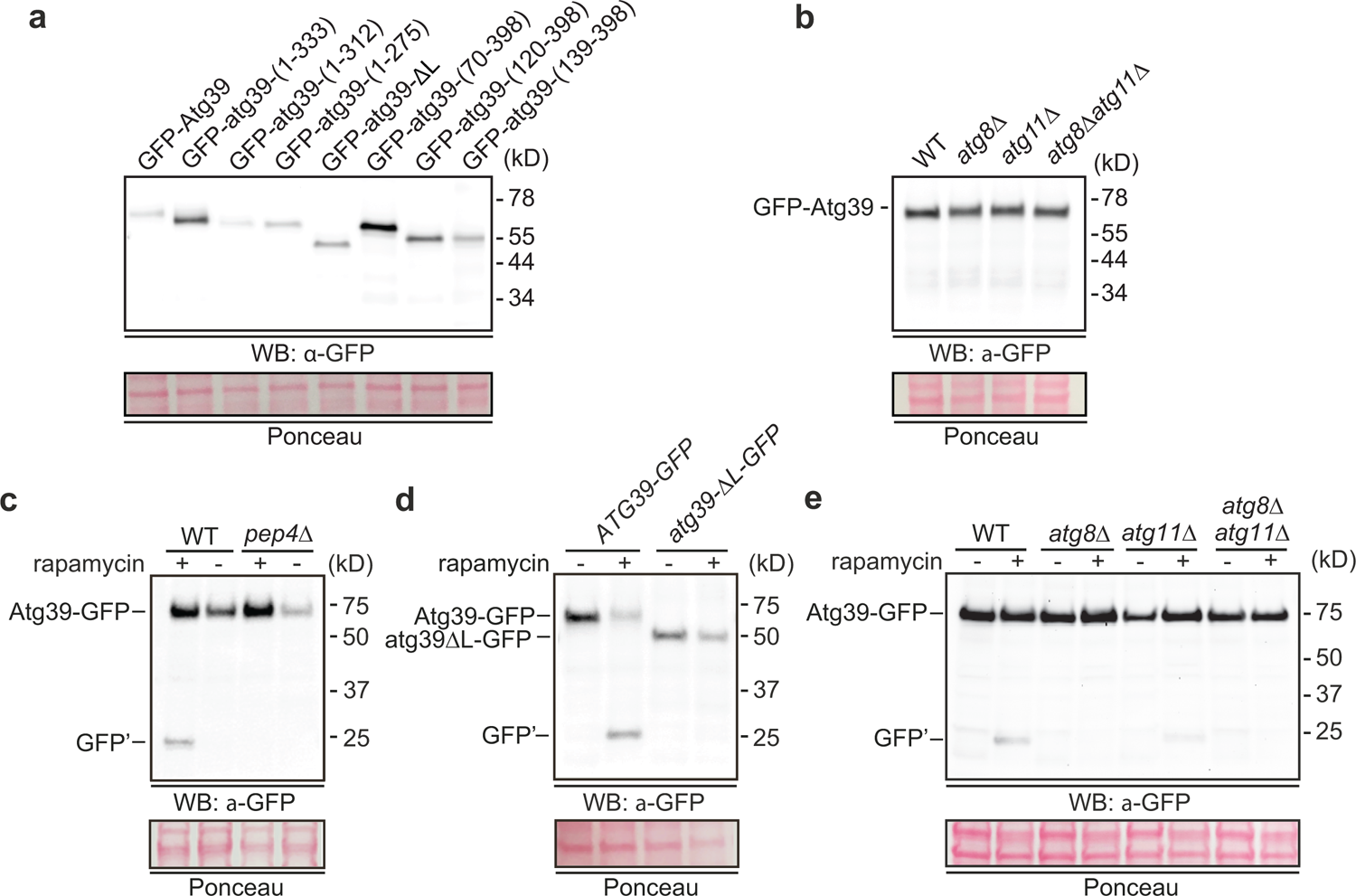
Assessment of total levels of Atg39 fusion proteins. **a-e**, Western blot of proteins from whole cell extracts derived from cells expressing the indicated GFP-fusions in indicated strains and drug treatments. GFP detected with anti-GFP antibodies (anti-GFP for a, b; anti-GFP.2 for c, e), HRP-conjugated secondary antibodies and ECL. Position of molecular weight standards (in kD) at right. To assess relative protein loading, a portion of the blots are shown stained with Ponceau.

**Extended Data Fig. 3.**
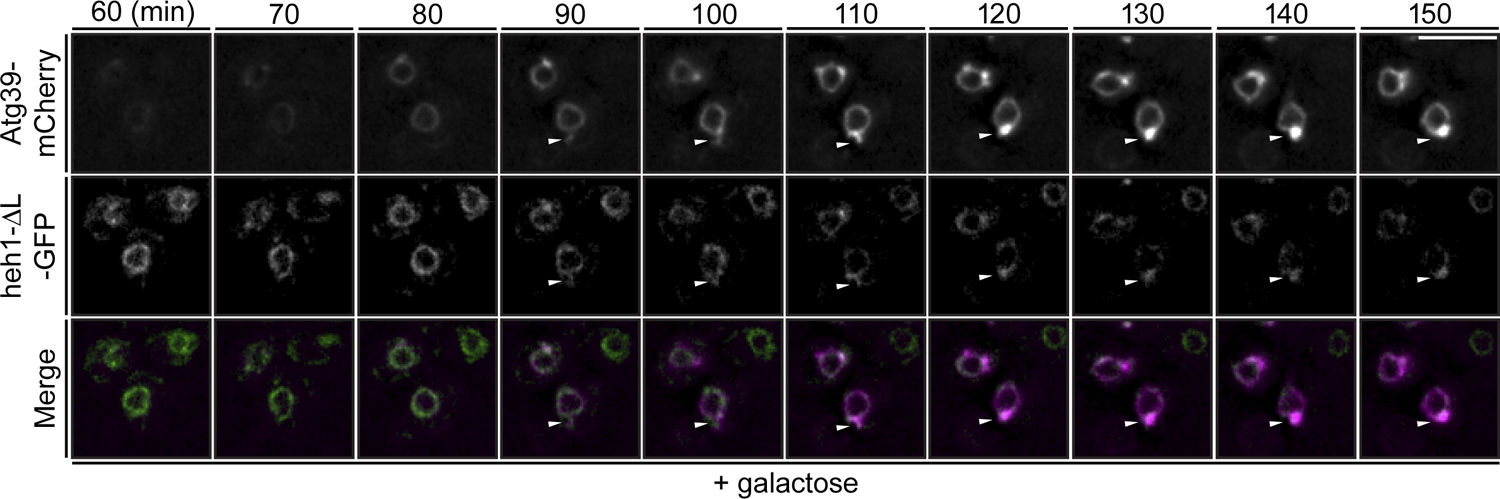
An Integral INM protein is enriched in NE blebs. **a**, Deconvolved fluorescence micrographs of a timelapse series showing cells expressing Atg39-mCherry (magenta) and heh1-ΔL-GFP (green) with merge at the indicated timepoints after addition of 2% galactose to induce Atg39-mCherry expression. White arrowheads indicate position of NE blebs. Scale bar is 5 µm.

**Extended Data Fig. 4.**
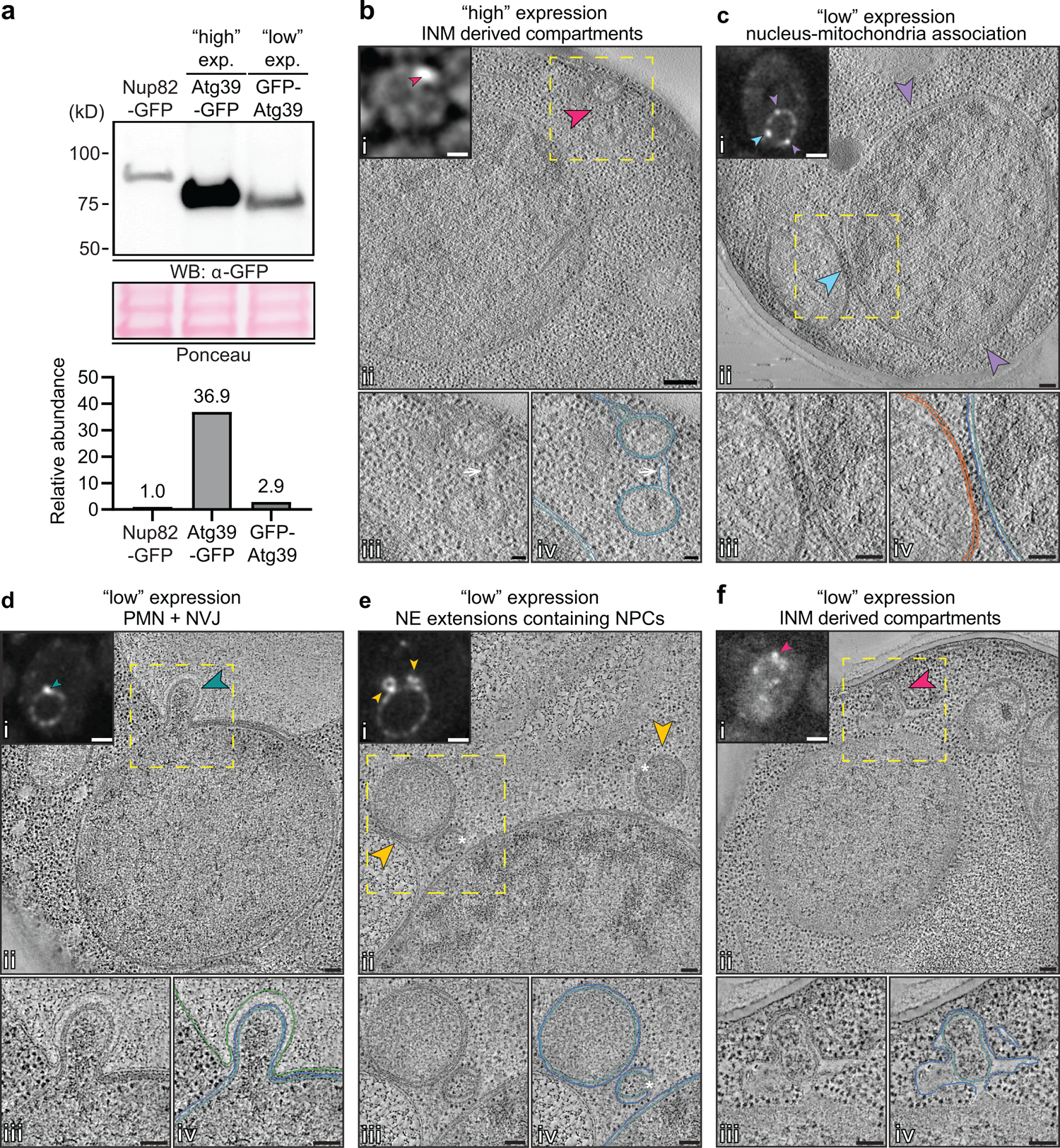
NE ultrastructure at sites of Atg39 accumulation. **a**, Western blot comparing relative levels of Atg39-GFP (“high” expression) and GFP-Atg39 (“low” expression) to endogenous levels of the nucleoporin, Nup82-GFP (∼1600 copies/cell). GFP detected with anti-GFP.1. Portion of blot stained with Ponceau shows relative protein loads. Bar graph is quantification by densitometry of the anti-GFP signal normalized to Nup82-GFP. **b-f**, CLEM tomograms from cells expressing “low” or “high” levels of Atg39 as indicated. (i) Fluorescence micrograph with arrows pointing to regions of interest similarly annotated in corresponding EM tomogram.(ii) Virtual slice from electron tomogram with the location of fluorescence from (i) indicated by similarly colored arrows. (iii) Magnification of boxed view in (ii). (iv) Annotation of virtual slices from panel (iii) with the ONM in blue, the INM in teal, vacuole in green and mitochondria in orange. Arrow points to continuity of ONM. Asterisks denote nuclear pores. Scale bars are 100 nm.

**Extended Data Fig. 5.**
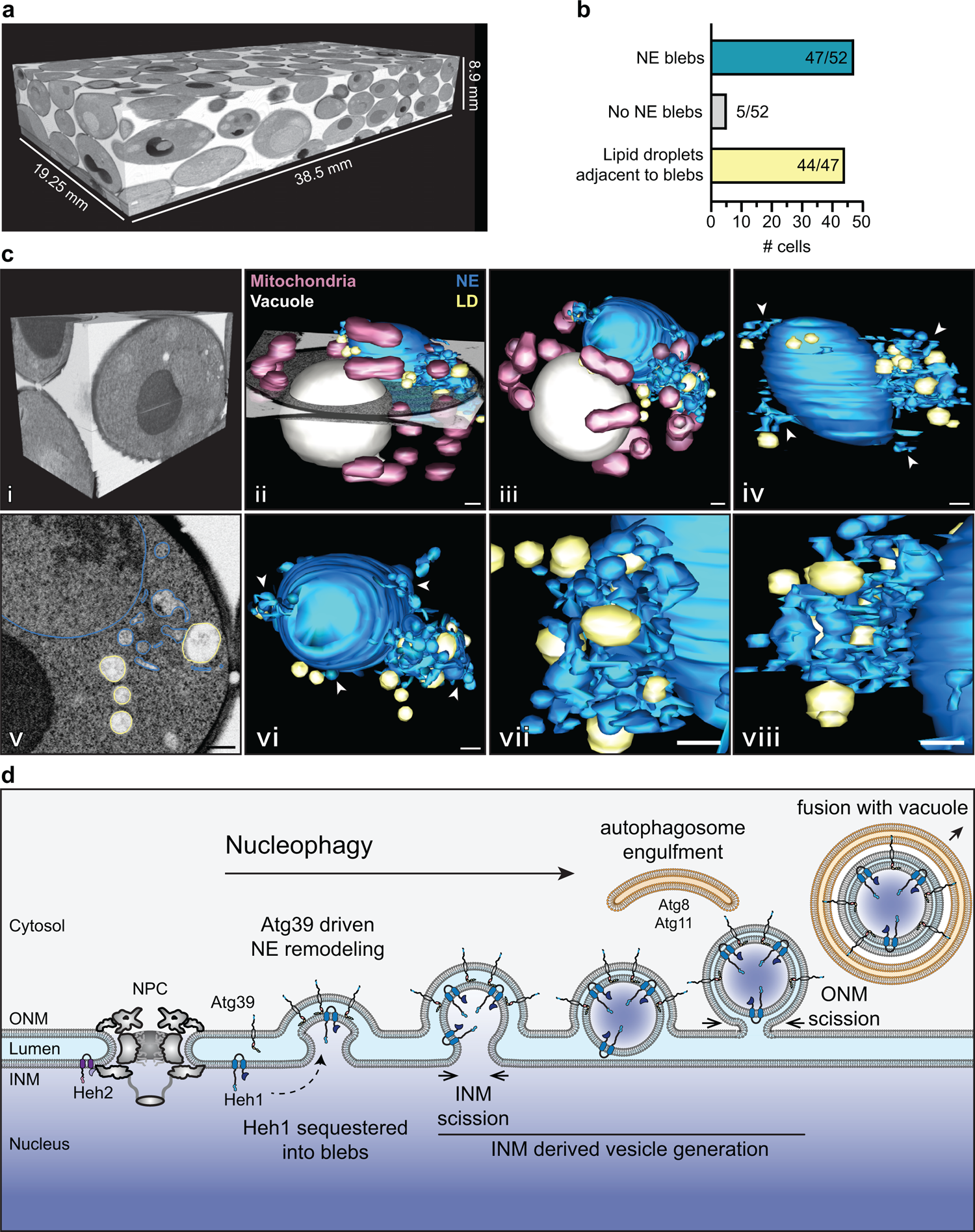
FIB-SEM of cells expressing Atg39 with model of nucleophagy. **a**, Total volume of FIB-SEM images of cells expressing Atg39-GFP. **b**, Bar chart of the quantification of NE blebs and associated lipid droplets observed in a total of 52 cells captured within the FIB-SEM volume shown in **a**. **c**, (i) Orthogonal view of a block of FIB-SEM images (from **a**) of a single cell with isotropic resolution of 7 x 7 x 7 nm voxels. (ii) surface rendering of NE (blue), lipid droplets (LDs, yellow), mitochondria (magenta), and vacuole (grey); an SEM image of a single Z-plane is also shown. (iii) top-down view of (ii). (iv) side view of surface rendering of just NE and LDs. Arrowheads point to NE blebs. (v) Annotated SEM image of a single z-slice with ONM blue, LD in yellow. (vi) top-down view of NE and LDs with arrows pointing to NE blebs. (vii, viii) Zoom-view and alternate angles of region containing NE blebs and LDs. Scale bars are 200 nm. **d**, A proposed outside-in model of nucleophagy. Atg39 localizes to the ONM and connects (directly or indirectly) to the INM through lumenal motifs. Evagination of the INM and selective capture of INM cargo (Heh1) requires Atg39. INM derived vesicles form after an INM scission event. Subsequent ONM scission would also be required to liberate the NE bleb before its capture by the autophagosome through interactions with the N-terminus of Atg39.

**Supplementary Table 1.**
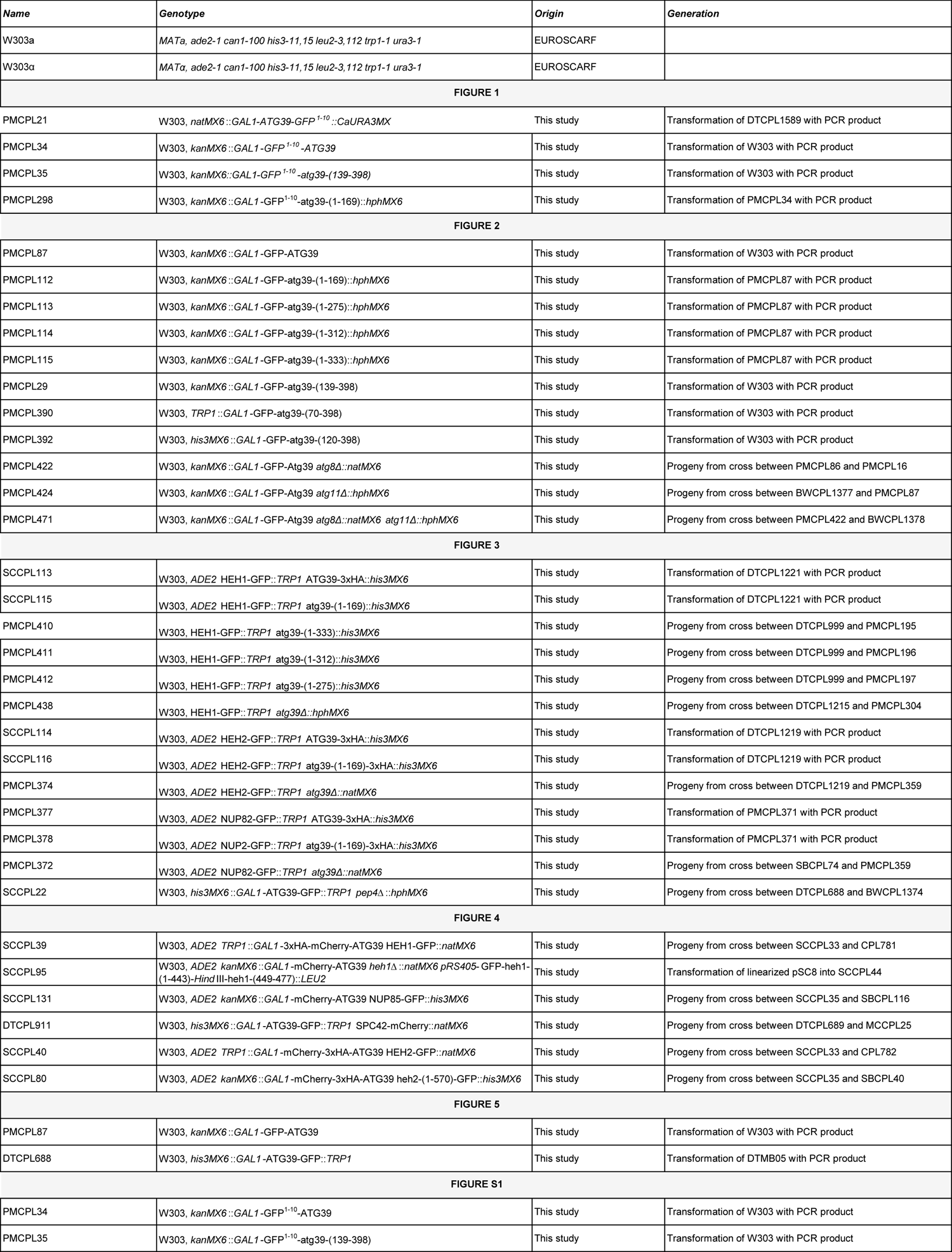
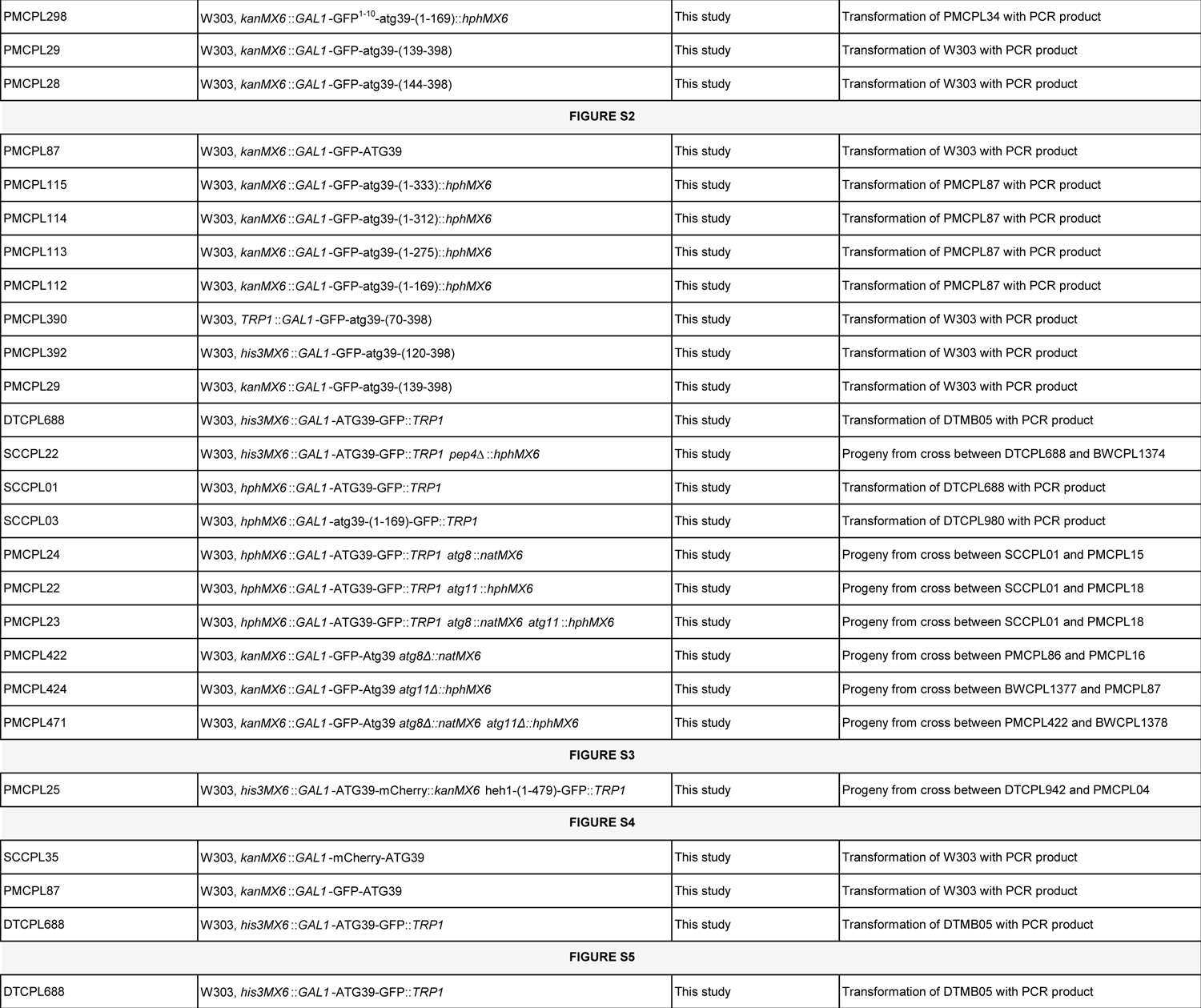
– *S. cerevisiae* strains used in this study.

**Supplementary Table 2.**
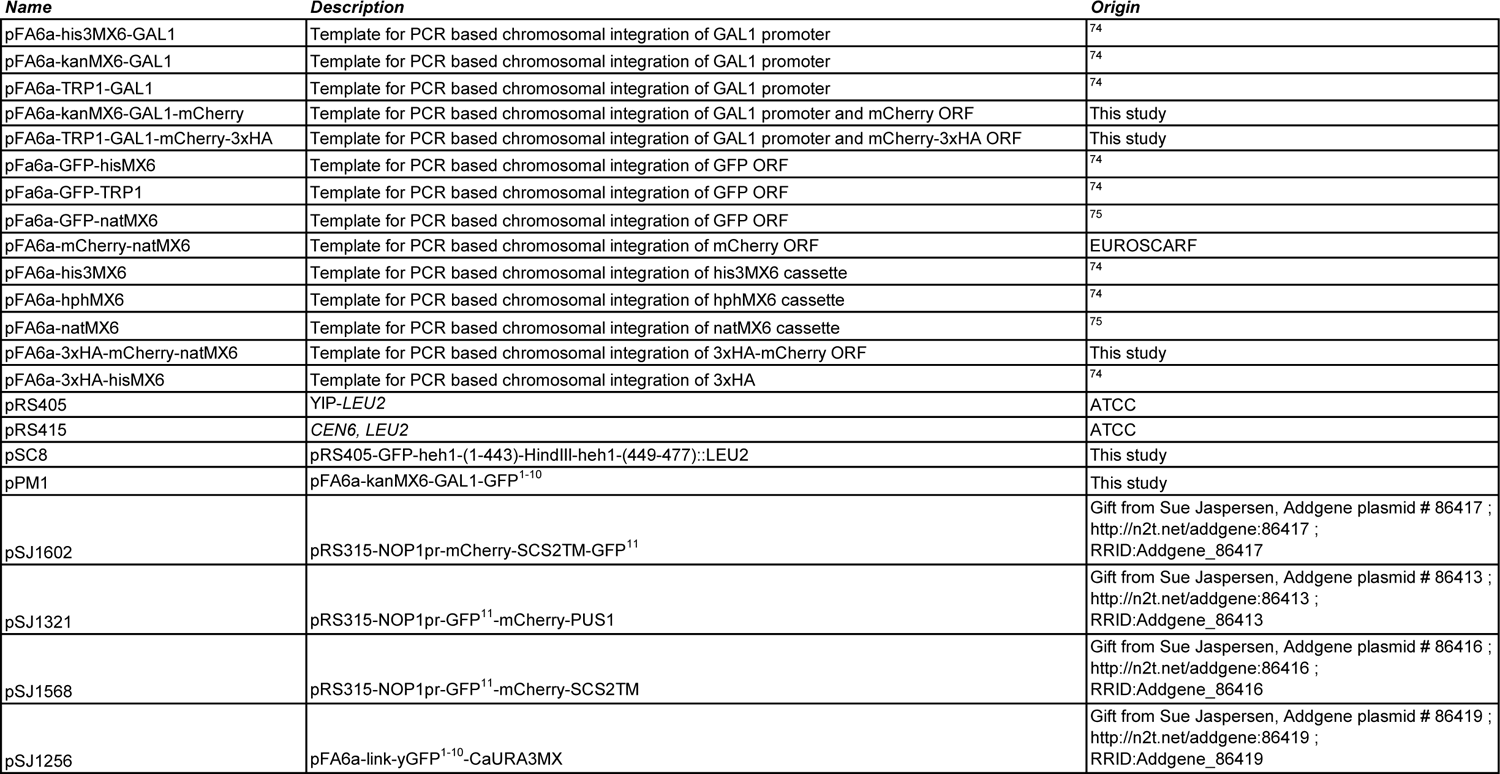
– Plasmids used in this study.

